# Assessing timing of fledging in a cavity-nesting passerine using temperature data loggers

**DOI:** 10.1101/2022.12.08.519594

**Authors:** Anna Dubiec, Tomasz D. Mazgajski

## Abstract

In altricial birds, the length of the nestling period, i.e. time from hatching until fledging (young leaving the nest) varies within and between species. In general, however, variation in the time of fledging and factors mediating such variation remain largely unexplored. To assess the time of nestlings leaving the nest, daily observer visits to the nest are usually done in the predicted fledging period. However, this might initiate premature fledging of young and/or increase the predation risk. The application of iButtons – coin-sized temperature data loggers, which are increasingly used in ornithological studies – may help to overcome these obstacles. We tested whether nest temperatures recorded with iButtons might be used to identify the date and hour of young fledging, i.e. when the last nestling in the brood left the nest, in a small cavity-nesting passerine – the Great Tit (*Parus major*). We installed iButtons in 38 nests when nestlings were 14-15 days old (hatching day = day 0) and verified the presence of nestlings during daily inspections starting on day 17 post-hatching or later. We found that the day of fledging could be accurately determined based on the difference between the temperature of the nest cup and the outside. The age of nestlings ranged between 17 and 22 days at fledging, with nearly 58% of broods fledging at the age of 20 and 21 days. The majority (81.6%) of broods fledged within 6 h after sunrise. We discuss the advantages and disadvantages of using iButtons to identify fledging time in altricial birds.

## 1. Introduction

Offspring of altricial species hatch undeveloped and, therefore, depend on parental care until they reach the stage of development that allows them to leave the nest. The length of period between hatching and fledging (leaving the nest) varies to a great extent among bird species (Cooney et al., 2020; Merrill et al., 2021; Remeš and Martin, 2002). In altricial, open-nesting species, like Eurasian Skylark (*Alauda arvensis*), the nestling period lasts for 7-11 days, while nestlings of other, often larger species, require more time to develop and leave the nests, even up to 58-64 days (White Stork (*Ciciona ciconia*)) (Billerman et al., 2022). The length of the nestling period varies also within species, both from the same or different populations (Johnson et al., 2004; Michaud and Leonard, 2000). For example, the length of the nestling period varied by nearly 100% (7-13 days) between the nests of the Horned Lark (*Eremophila alpestris*) in an alpine population (Zwaan et al. 2019). In general, however, the intra-specific variation in fledging age, and potential factors mediating such variation, remain largely unexplored (Moreno, 2020; Stodola et al., 2010; Yoda et al., 2017).

To accurately determine the age of nestlings leaving the nest, most commonly nests are daily visited around the expected fledging time (Bowers et al., 2013; e.g. Michaud and Leonard, 2000; Moreno, 2020; Stodola et al., 2010). However, the increased frequency of observer visits to the nests close to fledging time might initiate premature fledging (McCarty, 2001; Michaud and Leonard, 2000; Pietz et al., 2012), which may decrease the probability of post-fledging survival (but see Streby et al. 2013 for no or a positive effect of forced and premature fledging on fledgling survival of two songbird species). Moreover, frequent visits to the nests by observers might increase the risk of nest predation (Major, 1990), although the opposite or no effect has also been observed (Jacobson et al., 2011; Weidinger, 2008)

The application of temperature data loggers, which are increasingly commonly used in ornithological studies (e.g. Hartman and Oring, 2006), might help to overcome the obstacle of frequent nest visitation to determine the nestlings’ age at fledging. Temperature data loggers automatically measure and record temperature, according to the set frequency of data logging, for an extended time, limited by the memory capacity. To date, the temperature data loggers have been commonly used to examine the incubation behaviour (Arct et al., 2022; Ardia et al., 2009; Nord and Nilsson, 2012; Podlas and Richner, 2013), or to determine the events associated with temporary nest desertion or permanent termination of breeding attempts in response to predation or flooding (Arnold et al., 2006; Bayard and Elphick, 2011; Betuel et al., 2014; Hunter et al., 2016; Taylor, 2015). Other studies included an examination of air temperature within artificial and natural cavities (Ardia et al., 2006; Fairhurst et al., 2012; Maziarz et al., 2017; Pattinson et al., 2022) and an assessment of insulatory properties of bird nests (Deeming et al., 2020; Deeming and Campion, 2018).

Altricial nestlings are poikilothermic at hatching, but develop endothermy during the nestling period (Dunn, 1975). At the end of the nestling period, the body temperature of young is close to that of adult birds (at the active phase ca 41 °C) (Mertens, 1977; Prinzinger et al., 1991). Consequently, due to the endothermic activity of older nestlings, the temperatures of active nests are usually higher than in vacant nests, from which nestlings had fledged (Maziarz et al., 2020). As such, the nest temperatures could be a good indicator in monitoring the status of nests and the presence of live nestlings in them. Following the nestling departure, the nest temperatures should be similar to ambient temperatures, so the difference between internal (within a nest) and ambient temperatures would allow for pinpointing the date and hour of young fledging. Thus, collecting the data on nest temperatures would require only two visits to the nest: the first one – a few days before the expected fledging date to install a temperature data logger in the nest, and the second one – a few days later to check the content of the nest for the presence of nestlings and to retrieve the data logger if young absent. Despite the advantages of this method, such as low time investments and cost, the method was very rarely used specifically for assessing the time of young fledging in altricial birds (Ballance, 2018). Given the scarcity of the data, it is unclear to what extent the method could be successfully applied across different species.

We aimed to verify whether the fledging time could be accurately assessed by measuring the temperatures of bird nests with data loggers in a small cavity-nesting passerine – the Great Tit (*Parus major*). To establish the nest temperatures, we used iButton temperature data loggers (Maxim Integrated, USA; referred to hereafter as iButtons), which are stand-alone coin-sized loggers. By using the data collected with iButtons, we described the variation in fledging age and the diel pattern of fledging in the study species. Great Tit nestlings usually fledge between 16 and 22 days after their hatching, most of them in the mornings (Gosler et al., 2020; Radersma et al., 2015).

## 2. Materials and Methods

We collected data in a nestbox breeding population of Great Tits during April-June 2017 in the Sekocin Forest in central Poland, ca 10 km south-west of Warsaw (52°05′ N, 20°52′ E) (see Harnist et al. (2020) for a description of nestboxes and the study area). From early April we regularly checked nestboxes to record the first egg-laid date, clutch size, and hatching date (day = 0). On day 14-15 post-hatching, we ringed nestlings with a numbered aluminium ring, and placed one iButton in nesting material of 38 nests. All except five loggers were first mounted in a plastic iButton holder (DS9093S+, Maxim Integrated, USA; oval-shaped with dimensions of 46.9 mm × 30.9 mm) to help to immobilize the loggers within the nests. We positioned each iButton in a central part of the nest cup under ca 1-2 mm of the nesting material. Because the study aimed to verify whether fledging date and hour may be identified based on the comparison of nest and ambient temperatures, we deployed one logger (mounted in a plastic holder) to collect data on ambient temperature. We attached this logger to the outer side of the floor of the nestbox placed ca 3.5 m above the ground in the central part of the study area. We chose the position of the logger to keep it in shade during the daytime. We replaced the logger regularly to collect data on ambient temperature from the time we placed the first loggers in nests to the day the nestlings fledged from the last monitored nests. Only first broods were used in this study.

We used two models of iButtons: DS1921G-F5 and DS1922L (diameter: 17.35 mm, thickness: 5.89 mm). DS1921G-F5 measures temperature from -20 °C to +85 °C with an accuracy of 1 °C and resolution of 0.5 °C, and DS1922L – from -10 °C to +65 °C with an accuracy of 0.5 °C and resolution of 0.0625 °C. The use of two models was necessitated by the insufficient number of loggers of one type to record temperature in synchronized broods. The difference in the measurement accuracy between the two iButton models had no effect on identifying the date and hour of fledging, based on the comparison of the nest and ambient temperatures.

We programmed nest loggers and the logger recording ambient temperature to measure and record temperature synchronously (± 2 min) every 10 minutes. We chose the frequency of logging based on the maximum number of temperature readings that could be stored in a logger with a lower storing capacity (DS1921G-F5, up to 2048 readings) without the need for logger replacement, and considering the length of the nestling period of the study species.

Chosen logging frequency allowed to record temperature for 14 days, which corresponds to day 28 or 29 post-hatching depending on the day of logger installation. Based on literature data such a logging limit captures all fledging events in the study species (Gosler et al., 2020; Verhulst and Tinbergen, 1997). Loggers were programmed and temperature records were downloaded using OneWireViewer ver. 0.3.15.49 (Maxim Integrated Products, Sunnyvale, CA, USA).

To verify whether information on the nest activity status (active – nestlings present vs non-active – no live nestlings) may be reliably derived from the difference between the nest and ambient temperatures, the content of nest boxes was checked daily in the afternoon (after 1 pm) starting from day 17 post-hatching. Due to logistic constraints, in the subset of nests inspections started on day 18 post-hatching or later. The presence of nestlings was assessed through an entrance hole using a handheld flashlight with a flexible neck (Uni-max, Czech Republic) and a magnifying dental mirror. This method is used in studies on birds nesting in natural cavities (e.g. Wesolowski and Rowinski, 2014). The method minimizes the level of disturbance experienced by nestlings and prevents premature fledging. During nest-checks, we recorded the date, hour and outcome: nestlings present or absent. On the day we found that nestlings were absent, the logger was retrieved. We recorded the date and hour of logger retrieval, the number of dead nestlings, and in the subset of nests also crude estimation of the distance (in mm) between the bottom of the nest cup and the logger location. We collected data on dead nestlings, because their presence may affect temperature recorded by the logger in the nest. First, if some nestlings die before the last successful fledging event in a brood, their bodies, located at the bottom of the nest cup, may interfere with the transfer of the heat produced by live nestlings towards the logger. Second, when some nestlings die after the last successful fledging event in a brood, the moment when nest and ambient temperatures level out should be shifted in time, until the death of nestlings which failed to fledge. In both cases the drop in nest temperature following the last successful fledging event may be less distinct and consequently more difficult to detect than in nests without nestling mortality. We recorded the logger-bottom of the nest cup distance to verify whether the chosen method of logger installation (mounting in a plastic holder) successfully prevents extensive relocation of loggers in the nesting material.

### 2.1. Data analyses

To identify the time of the last successful fledging event in the nest, we compared the nest and ambient temperatures. As the time of fledging, we selected the hour corresponding to the start of a steady drop in the nest temperature until the nest and ambient temperatures leveled out. To compare nest and ambient temperatures, we plotted temperature records against time (Fig. 1, Hartman and Oring, 2006). Moreover, for each nest, we retrieved the minimum and mean difference between the nest and ambient temperatures over the period between midnight on the day the logger was placed in the nest and the hour identified as the moment of departure of the last nestling. The time of fledging was expressed as the number of hours since sunrise. Data on sunrise at the study site were obtained at https://www.timeanddate.com/. In the next step, we verified whether the date of fledging assessed with temperature data loggers corresponded with the outcome of the visual inspection of the nest activity status.

**Fig. 1.**
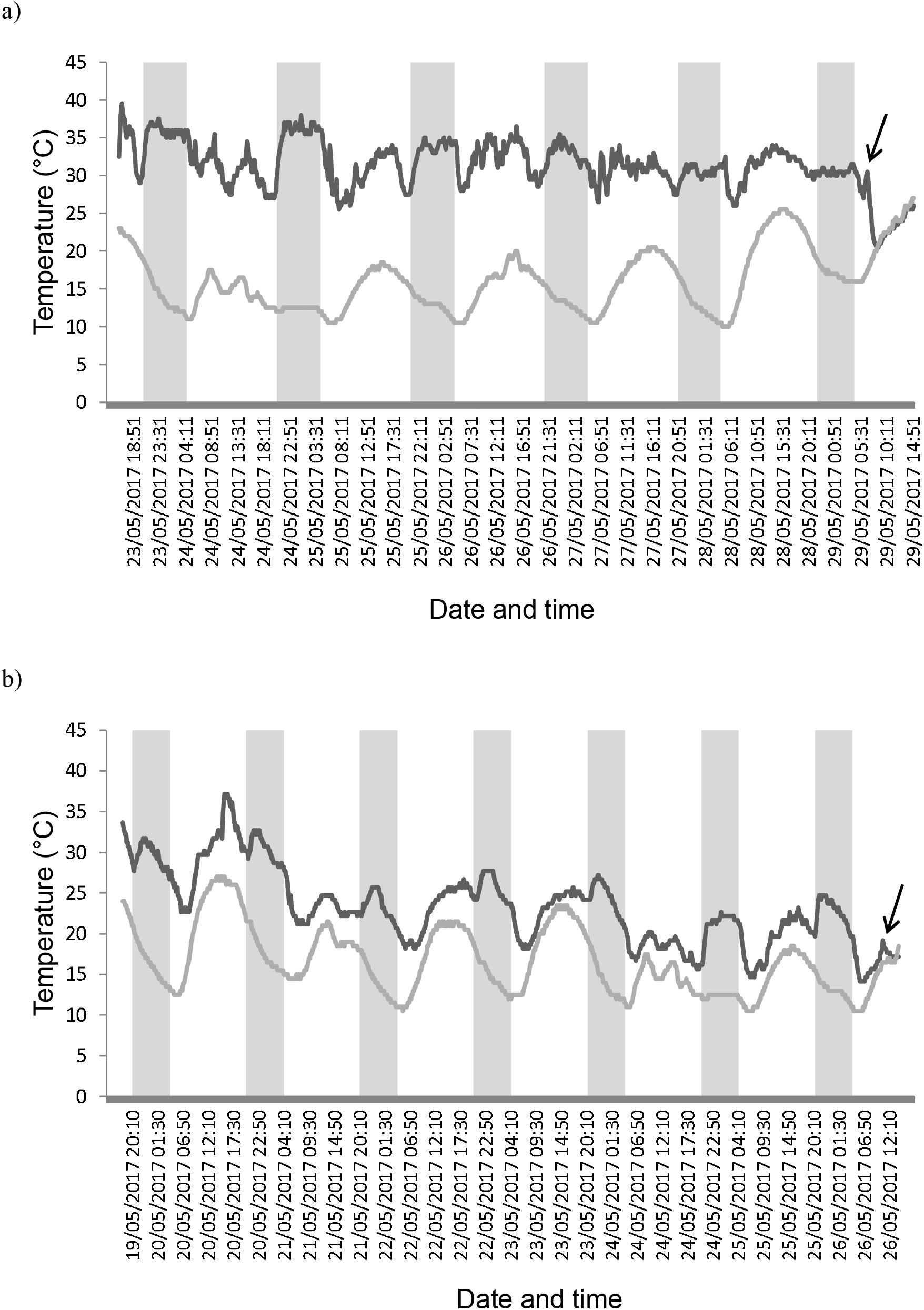
Records of the nest cup (black) and ambient (grey) temperatures between day 14 post-hatching (hatching day = day 0) and fledging day in two Great Tit nests in the Sekocin Forest in central Poland. Data for the nest with no (a) and partial (b) nestling mortality after day 14 post-hatching is presented. Nest cup and ambient temperatures were synchronously (± 2 min) recorded every 10 minutes. The arrow marks the time point of the presumed last successful fledging event in a brood. Periods of nighttime (between sunset and sunrise) are shaded.

Temperature records of the nest cup were available for 38 broods. However, in 7 broods inspection of the nestbox for the presence of nestlings was not carried out on daily bases or nestlings fledged before the first inspection. Since in such cases it was not possible to verify whether the temperature profiles of the nest cup corresponded with the presence of nestlings in the nestbox, such broods were excluded from the validation of the suitability of iButton temperature loggers for identifying the time of fledging. However, following the validation of the method, we used data from all 38 broods to assess fledging age and the diel pattern of fledging in the study population.

## 3. Results

### 3.1. Validation of the method

In all but one case, we retrieved the logger from the nesting material beneath the bottom of the nest cup. In one nest we found the logger on the rim of the nest, while the logger holder was in the nesting material. However, despite logger relocation, it was possible to identify fledging time based on recorded temperature. Some loggers (either those mounted in holders or placed in the nest without holders) changed their location relative to the bottom of the nest cup as the depth where they were found at the time of retrieval ranged from 0 to ca 20 mm.

In all 31 broods, the date of fledging could be accurately determined based on the comparison of the nest cup and ambient temperatures, as confirmed by the outcome of the visual inspection of nests for the presence of nestlings. This also applied to eight nests with dead nestlings (1-5 young over 14 days old) which were found on the day of the logger retrieval. In all nests, we identified the temperature record corresponding to the start of a gradual decrease in the nest temperature, which was followed by the nest and ambient temperatures leveling out (Fig. 1). On average, the minimum and mean per nest difference between the nest and ambient temperatures was 5.6 °C (range: 0.5–7.7 °C) and 12.4 °C (range: 6.2–17.7 °C), respectively.

### 3.2. Age and the diel pattern of fledging

Fledging of the entire broods was completed between days 17 and 22 post-hatching. Most broods fledged on days 20 and 21, 28.9% (n = 38) on each of these days. The average fledging age was 20.1 days (± 1.3 SD), while the median fledging age – 20 days (Fig. 2). The last nestlings in the brood departed from the nests between 26 minutes before and 14 h 18 minutes after sunrise, which corresponds to 4:01 am and 6:49 pm local time, respectively (Fig. 3). The mean and median time of fledging after sunrise were approximately 3 h 57 min and 2 h 46 min, which corresponds to 8:26 am and 7:15 am local time, respectively. The highest number of broods fledged between 2 h and 2 h 29 min after sunrise. Out of 38 monitored broods, 52.6% fledged within 3 h and 81.6% – within 6 h after sunrise.

**Fig. 2.**
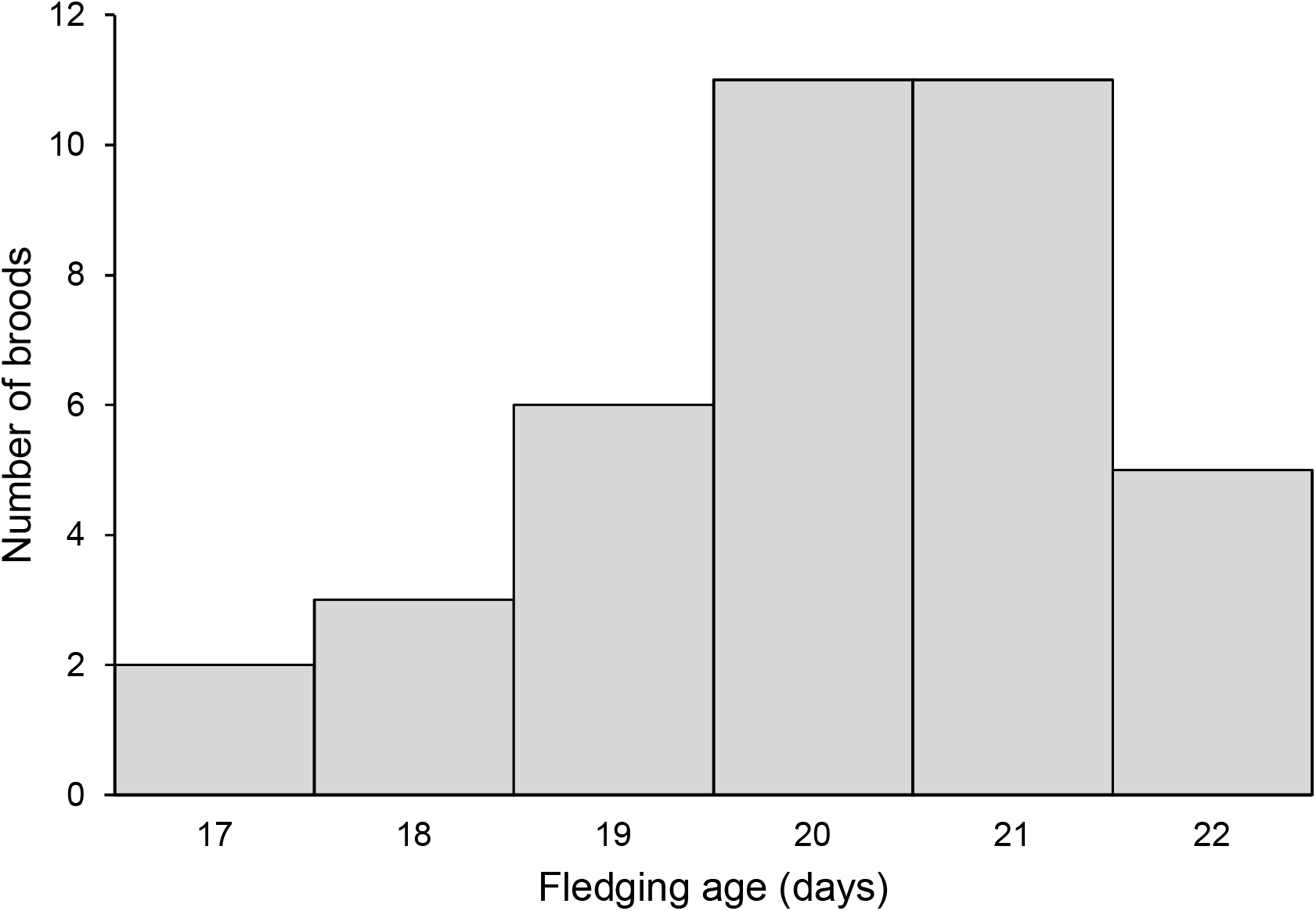
Frequency distribution of fledging age (in days) of Great Tit nestlings from first broods in the Sekocin Forest in central Poland in 2017. Fledging age was identified based on nest temperature records logged with iButtons.

**Fig. 3.**
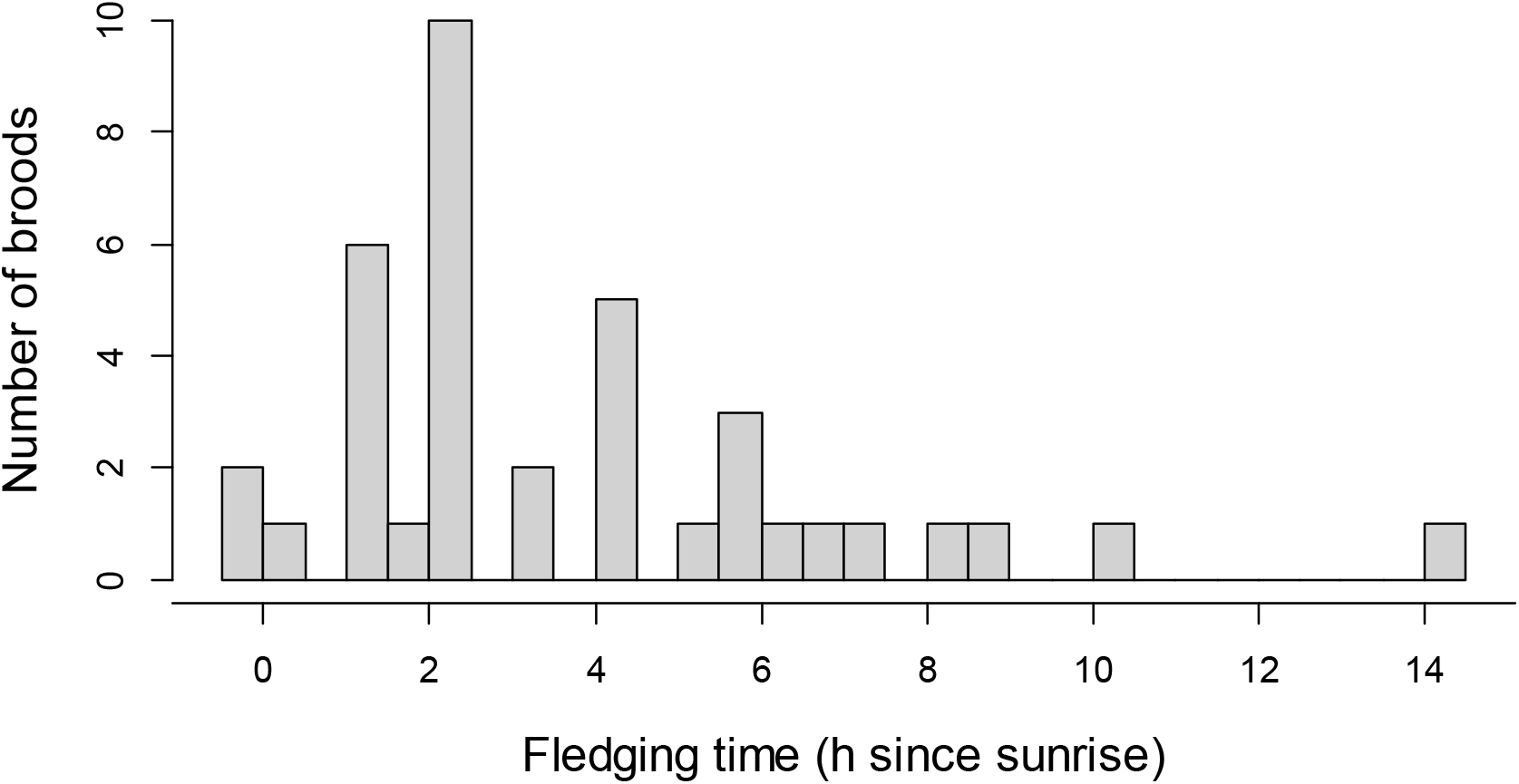
Frequency distribution of the time of fledging expressed as the difference in hours between sunrise and the moment of fledging of last nestling(s) in the nest.

## 4. Discussion

We show that iButton temperature data loggers may be successfully used to determine fledging age and the diel pattern of fledging in a cavity-nesting species that builds nests of dense structure (Gosler et al., 2020). In the study population of Great Tits, this method allowed for 100% accuracy in identifying the day when the last nestling(s) left the nest, as confirmed by daily nest visits around the predicted fledging day.

The two big advantages of this method over traditionally used frequent nest visitation are time and observer effort effectiveness and avoidance of disturbance of nestlings at the end of the nestling period when they might be provoked to leave the nest prematurely. Alternative remote methods such as radio-frequency identification technology (RFID) and video or trail cameras (Brandis et al., 2014; Iserbyt et al., 2018; Johnson et al., 2017, 2013, 2004; Pietz et al., 2012; Ribic et al., 2019; Surmacki and Podkowa, 2022) allow precise determination of timing of fledging, and, especially in the case of RFID technology, also fledging order (Radersma et al., 2015; Schlicht et al., 2012). However, given the much lower cost and easier installation, iButton temperature data loggers may be a method of choice in many studies focusing on the identification of fledging age in altricial birds.

When using iButtons to assess the time of fledging, one has to pay attention to the way the logger is installed in the nest. First, when unsecured, the logger may change its location in the nest. This may occur in response to nestling locomotor activity or when adult birds move the logger to other parts of the nest, e.g. to the nest rim (this study). With increasing distance from the nest cup containing nestlings, temperatures recorded by the logger should less accurately mirror temperatures of the nest cup. Such a reduction in temperature resolution in relation to the relocation of loggers in the nesting material has been previously observed in the Great Tit during the incubation stage when the main source of the heat is an incubating female (Smith et al., 2015). Second, iButton logger may be removed by the parent(s) outside of the nest. Smith (2011) reported frequent removal of unsecured iButtons during incubation by Great Tit females, while Nord and Nilsson (2012) reported such behaviour in incubating Pied Flycatchers (*Ficedula hypoleuca*), although in this species it was sporadic. To the best of our knowledge, no study so far looked into the frequency of iButton removal during different nesting stages. However, it is possible that such behaviour is more common during egg laying and incubation, when a logger may be perceived by parents as a parasitic egg, than during the nestling period. To reduce relocation and/or prevent removal of iButton loggers from the nest it is recommended that they are secured. This may be achieved by attaching strings/wire to the logger and wrapping them around the nesting substrate or e.g. a wooden stick placed beneath the nest (Schöll et al., 2020; Smith, 2011; Weintraub et al., 2016). In our study, we secured iButtons against relocation and removal by adult birds by mounting the logger in a plastic holder. Because almost all loggers were recovered not deeper than approximately 10 mm from the bottom of the nest cup, this method may be recommended in species building nests of dense structure. Moreover, only in one nest, the logger was removed by the adult bird from its location beneath the bottom of the nest cup by removing it from the plastic holder and placing it on the nest rim. By applying additional measures to secure the logger position in the holder, such a method should ensure full protection against relocation of loggers by parents in small bird species. In species that build nests of less dense structures and/or are larger, other methods may be preferred. For example, to monitor nest activity in the Tricolored Blackbird (*Agelaius tricolor*), an open-cup nester, that builds nests from leaves woven around plant stems, Weintraub et al. (2016) secured iButtons by sewing them into pieces of brown nylon stocking, which was then attached to green floral wire put through the bottom of the nest and wrapped around the nesting substrate.

While iButtons offer some advantages over other techniques of monitoring the nest activity (either remote or direct), this method has several limitations. First, with iButtons it is not possible to identify fledging age of individual nestlings. Temperature data loggers will only pinpoint the moment when the last nestling from the brood leaves the nest. In consequence, in multiparous species with brood fledging asynchrony i.e. when fledging of nestlings from the same brood is extended over time, this will result in the overestimation of the mean brood fledging age. However, the magnitude of such overestimation will depend on the degree of fledging asynchrony and the frequency of this phenomenon in the population. In species with narrow fledging asynchrony, i.e. when fledging of the entire brood spans across several hours but finishes on the same day, or is limited to a small proportion of broods in the population, the overestimation of the brood fledging age defined as the day when the last nestling left the nest, may be treated as negligible. In many songbirds all nestlings in the majority of broods complete fledging within 1 day, for example, in the Blue Tit (*Cyanistes caeruleus*) – in 55% (Schlicht et al., 2012), in the House Wren (*Troglodytes aedon*) – in 65% (Johnson et al., 2004) and in the Mountain Bluebird (*Sialia currucoides*) – in 83% of broods (Johnson et al., 2013). Also in the Great Tit in the majority of cases, the entire brood fledges on the same day (Verhulst and Tinbergen, 1997). Radersma et al. (2011) found that in this species fledging asynchrony within a brood ranged from 7 min 13 s to 2637 min (about 44 h) with an average of about 136 min. Second, with temperature records, it is not possible to distinguish between successful and depredated broods (either depredated fully or partially) if the predation event leaves no traces, such as feathers or disturbed nesting material (Weidinger, 2006). For example, Ball and Bayne (2012) monitoring 127 shrub–subcanopy nests representing 13 species found that among 78 nests that were classified as successful based on cues at the nest, 21 were in fact partially (14 nests) or entirely (7 nests) depredated based on video records.

And among 49 nests classified as depredated based on nest cues, in 5 nests 1 or more nestlings fledged. However, if nesting places are well secured against predators, for example, some types of nestboxes, the lack of live nestlings in the nest after the predicted fledging age may be relatively safely interpreted as a sign of successful fledging. For example, in the study population of Great Tits in the Sekocin Forest, no evident signs of nestling depredation were recorded over the whole period of population monitoring, i.e. for over 10 breeding seasons (own observation). Third, without video or trail camera records iButtons, similarly to fledging status assessed based on regular nest visits, do not allow to distinguish between unforced and forced fledging, either of the entire brood or some nestlings from the brood (Ball and Bayne, 2012). Fourth, iButtons may be used to correctly determine fledging time only under the condition that a difference between the nest and ambient temperatures is measurable. This may not be the case in regions with very high daytime ambient temperatures or when the nest is under direct sun exposure (Andersen and Freeman, 2022; Sutti and Strong, 2014). For example, Andersen and Freeman (2022) showed that in the Botteri’s Sparrow (*Peucaea botterii*), a species that nests in hot, semiarid grasslands, during the hot period of the day the cessation of nest activity was correctly identified only in 46% of nests, while during the cooler period of the day (when nest temperature was on average 3.9 °C higher than ambient temperature), termination of the nest activity was correctly assigned in all nests. However, even though the hour of termination of the nest activity may not always be correctly assigned, a distinct difference between nest and ambient temperatures during the cooliest parts of the day (night, early morning, late evening) should still allow for correct identification of the day of fledging.

The average fledging age in the study population was 20.1 days and in the broods with the longest nestling period nestlings departed from the nest 5 days later than in nests with the shortest nestling period. Great Tit nestlings in central Poland fledged approximately a day later than in the populations on the island of Vlieland (the Netherlands) and near Bern (Switzerland) and half a day later than in the population in Lauwersmeer in the north of the Netherlands (Basso and Richner, 2015; Radersma et al., 2011; Verhulst and Tinbergen, 1997). However, in order not to introduce any bias in the comparison of fledging ages among populations, data should be preferably collected with the use of the same method and use the same definition of fledging age. For example, while in the studies in the Sekocin Forest and Vlieland populations fledging age was defined as the day when the last nestlings left the nest, the study in the Lauwsmeer population, which applied RFID technology, defined fledging age at the brood level as the average of fledging age of individual nestlings. Because of that in broods with extended fledging asynchrony, such an approach introduces an underestimation in fledging age when compared to this parameter expressed as the day when the last nestling left the nest. The majority of Great Tit broods fledged early in the morning with 52.6% of broods fledging within 3 h and 81.6% – within 6 h after sunrise. Fledging at this time of the day is typical for the Great Tit (Lemel, 1989; Radersma et al., 2015; Verhulst and Hut, 1996) as well as for other passerines and is being associated with the developmental stage of nestlings and predation risk (Chiavacci et al., 2015; Santema et al., 2021 and references therein; Schlicht et al., 2012).

Summing up, this study shows that iButton temperature data loggers may be used to correctly determine fledging age as well as the hour of fledging in an altricial cavity-nesting species. To increase the accuracy of assessment, iButtons are recommended for populations with low predation risk. In general, it is also recommended that iButtons are secured in the nest to prevent their removal and/or relocation within the nest by adult birds. Using iButtons, we showed that in the study population there is substantial variation in the age when nestlings leave the nest, which may potentially translate into differences in such fitness-related traits as survival.

## Acknowledgements

We thank the Forest Research Institute in Sękocin Stary for logistic support and Marta Maziarz for comments that improved the manuscript. This study was partly supported by the Polish Ministry of Science and Higher Education/National Science Centre grant to TDM (grant no. N304 345139).

## Conflict of interest

Authors declare that they have no conflict of interest.

## Ethics Statement

All applicable national and institutional guidelines for the care and use of animals were followed. The procedures were carried out under the permit issued by the Regional Directorate for Environmental Protection in Warsaw, Poland (permit no. WPN-I.6401.159.2017.LM).

## Notes

### Competing Interest Statement

The authors have declared no competing interest.

